# L-cysteine inhibited the growth of *Vibrio parahaemolyticus* via increasing the ROS level

**DOI:** 10.64898/2026.01.16.699907

**Authors:** Qiuyan Yang, Huaipeng Fang, Liting Xu, Min Meng, Qingxi Han, Weiwei Zhang

## Abstract

The emergence of multidrug-resistant *Vibrio parahaemolyticus* poses a severe threat to mariculture sustainability, highlighting the urgent need for eco-friendly antimicrobial agents. In this study, we demonstrated that L-Cysteine (L-Cys) functions as an inhibitor of *V. parahaemolyticus*, with a minimum inhibitory concentration of 7.5 mM. Microscopic observation and viability assays revealed that L-Cys compromises bacterial membrane integrity, ultimately leading to cell death. Further investigation indicated that the antibacterial effect is primarily attributed to the intracellular production of hydrogen sulfide (H_2_S) generated by L-Cys metabolism. Transcriptomic and biochemical analyses showed that L-Cys induced metabolic reprogramming by suppressing fatty acid β-oxidation, one-carbon metabolism, and antioxidant enzymes. This disruption of redox homeostasis results in accelerated accumulation of reactive oxygen species (ROS). In addition to its antibacterial effect, L-Cys also effectively reduced the virulence factor of bacterial motility. Finally, L-Cys demonstrated broad-spectrum antimicrobial activity against other pathogenic *Vibrio* species, including *V. alginolyticus* and *V. anguillarum*. Our findings suggest that L-Cys is a promising antimicrobial agent inducing ROS to mediate membrane disruption, with the advantages of cost-effectiveness and environmental safety for controlling vibriosis.

**Importance:** *Vibrio parahaemolyticus* is a significant pathogen in aquaculture and a common cause of seafood-borne gastroenteritis worldwide. The increasing prevalence of antibiotic-resistant strains of this bacterium highlights the need for alternative control agents. This study shows that L-cysteine (L-Cys) inhibits the growth of *V. parahaemolyticus*. Our data indicate that L-Cys is metabolized to produce hydrogen sulfide, which contributes to the accumulation of ROS and disrupts bacterial membrane integrity. Additionally, L-Cys reduces bacterial motility and shows inhibitory effects against other *Vibrio* species. These findings suggest that L-Cys may represent a useful agent for managing *Vibrio* infections in aquaculture settings.

## Introduction

Bacteria of the genus *Vibrio* are widely distributed in marine and estuarine environments (1). They can cause vibriosis, posing serious threats to aquaculture, with pathogenic isolates such as *Vibrio parahaemolyticus*, *V. alginolyticus*, *V. harveyi*, *V. anguillarum*, *V. crassostreae*, *V. splendidus*, and *V. mediterranei* frequently reported in farmed aquatic species (1, 2). These pathogens, particularly *V. parahaemolyticus*, not only lead to disease and mortality in farmed organisms, but also cause human foodborne illnesses such as acute gastroenteritis and septicemia through the food chain, thereby posing significant risks to public health (1, 3).

*V. parahaemolyticus* is a halophilic Gram-negative bacterium and a typical aquatic-borne pathogen, attracting considerable concern due to its ecological adaptability and strong pathogenicity (4). It infects a wide range of aquatic animals, including nearly all major aquaculture species such as shrimp, crabs, fish, mollusks, and echinoderms (4). Moreover, it is the leading cause of seafood-related foodborne illnesses worldwide, accounting for a large proportion of food poisoning cases in coastal regions, with infection rates increasing annually(5, 6). Its virulence is mediated by multiple factors, including thermostable direct hemolysin (TDH) and TDH-related hemolysin (TRH), which disrupt host cell membrane integrity (7, 8); the Type III Secretion System (T3SS), which drives cytotoxicity and intestinal pathogenesis (8, 9); and the Type VI Secretion System (T6SS) subtype T6SS2, which promotes bacterial adhesion to host cells (10). These virulence mechanisms contribute to severe diseases and substantial economic losses in aquaculture. For example, *V. parahaemolyticus* infection in shrimp *Penaeus vannamei* leads to Acute Hepatopancreatic Necrosis Disease (AHPND), causing mortality rates exceeding 80% and resulting in significant economic damage (11, 12). In shrimp *Penaeus monodon*, the secretion of metalloproteases and serine proteases by *V. parahaemolyticus* degrades hepatopancreatic tissues, causing necrosis and exhibiting high toxicity (13). In recent years, this pathogen has also been identified as the primary cause of translucent post-larvae disease (TPD) in *P. vannamei* post-larvae, leading to high mortality through colonization of the digestive tract and severe necrosis of hepatopancreatic and midgut epithelial cells, which results in the characteristic translucent appearance of affected larvae (14).

Controlling vibriosis outbreaks in aquaculture remains challenging. On one hand, elevated water temperatures (30-37°C) and increased organic matter promote *Vibrio* proliferation (15, 16). On the other hand, indiscriminate antibiotic use has exacerbated antimicrobial resistance (17). *V. parahaemolyticus* strains exhibit resistance to multiple antibiotics, including β-lactams and sulfonamides (17, 18). Consequently, eco-friendly agents and strategies that can control *Vibrio* populations without inducing resistance have gained attention. These include the use of probiotics such as *Bacillus* spp. and lactic acid bacteria (19), bacteriophages (20), natural plant extracts like tea polyphenols (21), and amino acid-based antimicrobials such as ε-polylysine and D-amino acid peptides (22, 23). These strategies operate through distinct mechanisms: probiotics produce bacteriocins or organic acids (19); bacteriophages lyse bacteria by recognizing surface receptors and injecting genomic DNA (20); tea polyphenols disrupt cell membranes and inhibit key metabolic enzymes (21); ε-polylysine forms pores in the cell membrane, causing leakage of intracellular components (22); and D-amino acid peptides interfere with peptidoglycan synthesis and biofilm formation (23). Notably, amino acid-based antibacterials offer advantages such as environmental safety, cost-effectiveness, and low resistance risks (22).

Recently, L-cysteine (L-Cys) has attracted attention for its antibacterial activity, showing inhibitory effects against *Escherichia coli*, *Staphylococcus aureus*, *Listeria monocytogenes*, and *Salmonella enteritidis*, with the strongest activity observed against *S. aureus* (24). At a concentration of 1000 mg/L, L-Cys significantly delays spore germination and inhibits mycelial growth of the fungus *Monilinia fructicola*, a pathogen responsible for postharvest peach brown rot (25). In *Vibrio* spp., hydrogen sulfide (H_2_S) derived from intracellular L-Cys metabolism via cystathionine β-synthase (CBS) enhances *V. cholerae* colonization by promoting iron-dependent catalase KatB activity (26). However, the effect of L-Cys on the important pathogenic *Vibrio* species in aquaculture remains unexplored.

This study aims to systematically evaluate the inhibitory effect of L-Cys on *V. parahaemolyticus*. The underlying mechanism were investigated through transcriptomic analysis, combined with assessments of cell membrane integrity, hydrogen sulfide (H_2_S) production, antioxidant enzyme activity, and intracellular reactive oxygen species (ROS) levels. Furthermore, the impact of L-Cys on virulence factors of *V. parahaemolyticus* was examined. Finally, the broad-spectrum inhibitory activity of L-Cys against other pathogenic *Vibrio* species was assessed. This study not only elucidates how L-Cys inhibits the growth and virulence of *V. parahaemolyticus* but also provides a potential tool for controlling *Vibrio* infections in aquaculture.

## Materials and Methods

### Bacterial Strain and Culture

The *V. parahaemolyticus* strain YDE17 used in this study was obtained from our laboratory collection. The strain was routinely cultured in 2216E medium at 28 °C. For carbon source screening experiments, the following media were prepared: M9 medium (Haibo, Qingdao, China) without D-glucose (Coolaber, Beijing, China); M9 minimal medium supplemented with 0.4% D-glucose as carbon source; and M9 agar plates containing 1.2% agar. All media were adjusted to 3% (*w*/v) salinity by adding NaCl. In subsequent liquid culture experiments, bacteria were inoculated at 1:100 (*v*/v) and incubated at 28 °C with shaking at 160 rpm.

### Screening of Amino Acids as the Sole Carbon Sources

The selection of amino acids used by *V. parahaemolyticus* YDE17 was performed following the method described by Zhang et al. (27), with minor modifications. Twenty L-amino acids (Coolaber, Beijing, China) were individually supplemented as the sole carbon source in M9 agar plates. Briefly, 1 mL aliquots of bacterial culture at OD_600_ ≈ 0.5 were collected with three biological repeats, washed three times with M9 medium, and resuspended in the same medium. For liquid assays, the washed cells were inoculated into M9 medium supplemented with 40 mM of each L-amino acid. Cultures were incubated at 28 °C with shaking at 160 rpm in a constant-temperature shaker (Jiecheng, Shanghai, China) for 12 h, and OD_600_ was measured using a microplate reader (FlexA-200, Allsheng, Hangzhou, China). For plate assays, the bacterial suspensions were 10³-fold diluted, and 100 μL of each dilution was spread onto M9 agar plates containing the corresponding L-amino acid, with three technical replicates per sample. After incubation at 28 °C for 24 h, single colonies were enumerated.

### The inhibitory effect of L-Cys on the growth of *V. parahaemolyticus* YDE17

The inhibitory effect of L-Cys was evaluated according to the method described by Wang et al. (24). Briefly, *V. parahaemolyticus* YDE17 was cultured to exponential phase in M9 minimal medium. Cells from three independent cultures were harvested, washed three times with M9 medium, and resuspended to an OD_600_ of 0.5. Resuspended cells were then inoculated into M9 minimal medium supplemented with L-Cys at concentrations of 0, 1.25, 2.5, 5, 7.5, 10, 20, 30, 40, and 50 mM, with three biological replicates per concentration. A control group without L-Cys was included. All 90 cultures were incubated at 28 °C for 24 h, after which the OD_600_ of each culture was measured.

To assess growth in 2216E medium, 50 μL aliquots of *V. parahaemolyticus* YDE17 culture were inoculated into fresh 2216E media containing L-Cys concentrations ranging from 0 to 50 mM, with three biological repeats were performed. Cultures were incubated at 28 °C for 18 h, with OD_600_ measured every 2 h using a microplate reader.

### Scanning Electron Microscopy

Samples for scanning electron microscopy (SEM) were prepared following the method of Ishrat et al. (28). *V. parahaemolyticus* YDE17 was cultured in fresh 2216E medium, with or without 5 mM L-Cys, and incubated at 28 °C with shaking at 160 rpm until the exponential phase was reached. Cells were harvested by centrifugation at 3000 × g for 10 min, fixed with 3% glutaraldehyde at 4 °C for 2 h, and washed three times with 0.1 M PBS. Subsequently, samples were dehydrated through a graded ethanol series (30%–100%), subjected to critical-point drying, sputter-coated with gold, and examined using a Hitachi S-3400H scanning electron microscope (Hitachi, Japan).

### Membrane Integrity Assays

Membrane integrity was evaluated using the LIVE/DEAD Bacterial Viability Kit (Beyotime Biotechnology, Shanghai, China) according to the manufacturer’s instructions. The kit utilizes the nucleic acid stains N,N-dimethylaniline N-oxide (DMAO), a membrane-permeable green fluorescent dye that labels all bacterial cells, and propidium iodide (PI) that only enters cells with compromised membranes and fluoresces red. *V. parahaemolyticus* YDE17 in the exponential growth phase was harvested, resuspended in sterilized seawater, and divided into four aliquots. Three aliquots were treated with 5 mM L-Cys at 28 °C for 30 min, then stained with the working dye solution and incubated in the dark at 37 °C for 15 min. The remaining aliquot, which received no L-Cys treatment, served as the control. A 10 μL sample of each bacterial suspension was placed on a slide and examined under an inverted fluorescence microscope (Nikon Eclipse TI-S, Japan). Cells with intact membranes displayed green fluorescence of DMAO, while those with damaged membranes exhibited red fluorescence of PI.

### H_2_S Measurement

The Lead Acetate Paper Test was conducted following the method of Ma et al. (26). Five milliliters of fresh 2216E medium was aliquoted into 15 mL glass flasks, each supplemented with L-Cys at final concentrations of 0, 0.125, 0.25, 0.5, 1, 1.25, 2.5, and 5 mM, respectively. Three independent replicates were prepared per concentration, totaling 24 flasks. Each flask was inoculated with 1% (*v*/v) bacterial culture. A sterilized moistened lead acetate test strip (Sssreagent, Shanghai, China) was fixed to the mouth of each flask using a rubber stopper. Following incubation at 28 °C with shaking at 160 rpm for 12 h, color changes on the test strips were recorded.

In a parallel experiment, 24 mL of inoculated 2216E medium was divided into 12 aliquots, supplemented with L-Cys at concentrations of 0, 1.25, 2.5, and 5 mM, with three biological replicates per concentration. After incubation under the above conditions for 12 h, the concentration of H_2_S gas in each tube was measured using an EDKORS portable H_2_S gas detector (Changzhou, China).

### Bacterial motility Assay

Swimming and swarming assays were performed according to the method of Li et al. (29). 2216E agar plates containing 0.25% (*w*/v) agar (for swimming) and 0.6% (*w*/v) agar (for swarming) were prepared with or without 10 mM L-Cys. For the swimming assay, 2 μL of *V. parahaemolyticus* YDE17 cells were inoculated into the center of the 0.25% agar plate. For the swarming assay, 5 μL of bacterial suspension were spotted onto the surface of the 0.6% agar plate. In each assay, three independent cultures adjusted to an OD_600_ of 0.5 were used. After static incubation at 28 °C for 12 h, the radius of bacterial migration was measured.

### Real-time reverse transcriptase PCR (RT-PCR)

*V. parahaemolyticus* YDE17 was inoculated into fresh 2216E medium supplemented with 5 mM L-Cys and incubated at 28 °C with shaking at 160 rpm for 24 h. A control group was cultured in parallel using 2216E medium without L-Cys. Both conditions were performed in triplicate. Bacterial cells were harvested and washed with sterilized seawater to an OD_600_ of approximately 0.5. Total RNA was extracted using the MiniBEST Universal RNA Extraction Kit (TaKaRa, Beijing, China). Subsequently, RNA was reverse-transcribed and purified using the EasyScript® All-in-One First-Strand cDNA Synthesis SuperMix for qPCR (One-Step gDNA Removal; TransGen Biotech, Beijing, China). Quantitative real-time RT-PCR was carried out on an ABI 7500 Real-Time Detection System (Applied Biosystems, USA) under the following cycling conditions: 94 °C for 30 s, followed by 40 cycles of 94 °C for 5 s and 60 °C for 30 s. Primer sequences are listed in Table 1. For each gene, six technical replicates were included in a 96-well plate. The amplification efficiencies of all primers were validated and fell within the acceptable range of 90–110%. Data are presented as mean ± standard deviation (SD).

**Table 1.**
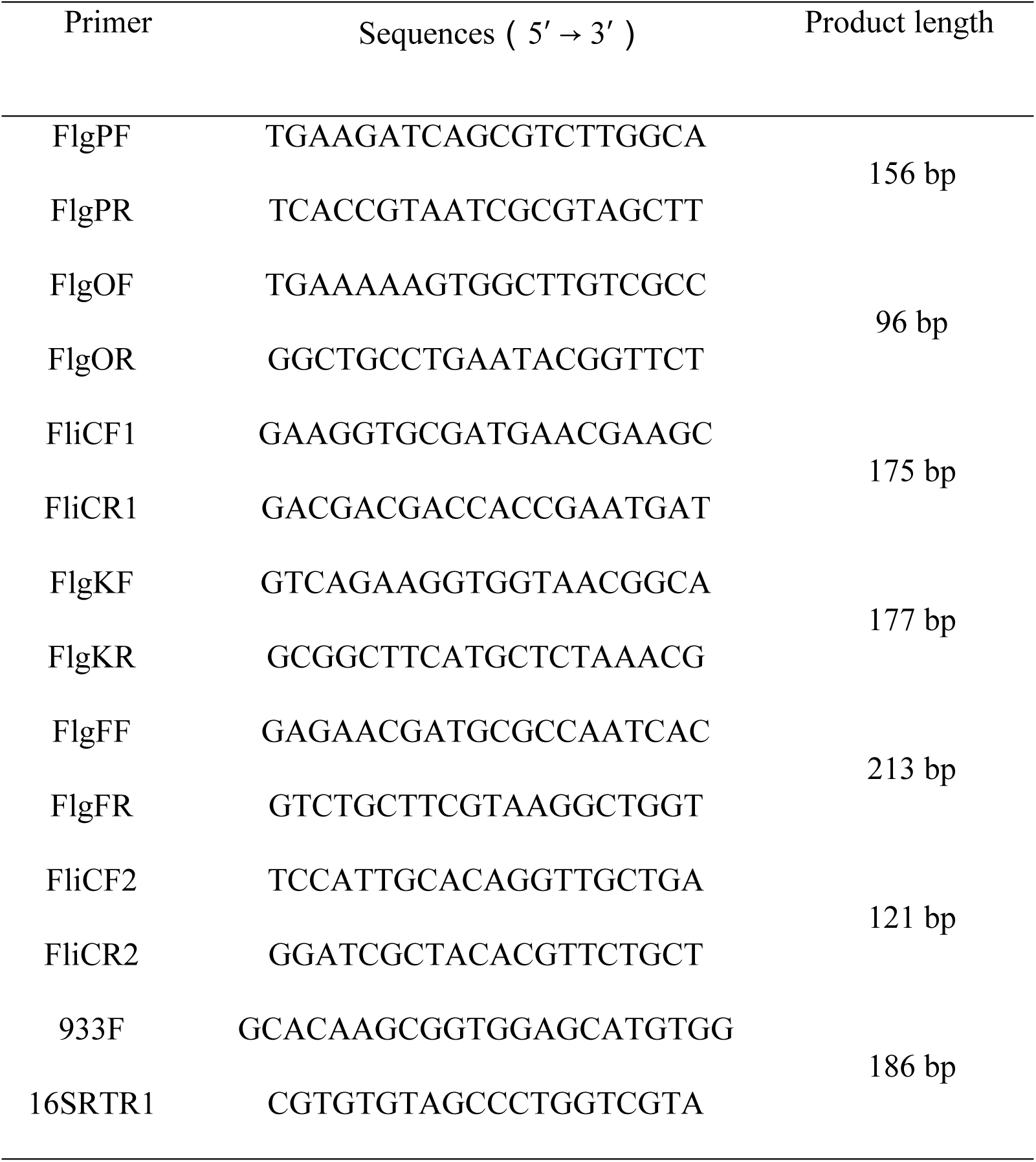
Primers used for quantitative real-time RT-PCR.

### Transcriptomic Sequencing and Analysis

*V. parahaemolyticus* YDE17 was cultured in 2216E medium for 24 h, with or without 5 mM L-Cys. Cells were harvested and adjusted to an OD_600_ of 0.5. RNA extraction, cDNA library construction, and sequencing were performed following the method of Dmitri et al. (31). Total RNA was extracted using TRIzol Reagent (Thermo Fisher Scientific, USA), and RNA integrity was assessed on an Agilent 2100 Bioanalyzer (Agilent Technologies, USA). Following rRNA depletion, mRNA was purified, fragmented, and used for cDNA library construction. Libraries were prepared and sequenced on the Illumina HiSeq/MiSeq platform by Novogene (Beijing, China). The *V. parahaemolyticus* reference genome ASM19609v1 (GCF_000196095.1) was used for read alignment. Gene expression levels were quantified in FPKM using RSEM, and differentially expressed genes (DEGs) were identified with a significance threshold of padj < 0.05. GO and KEGG enrichment analyses were performed using the GOseqR package and KOBAS software, respectively (Novogene, Beijing, China).

### ROS Measurement

Intracellular ROS levels were measured using the fluorescent probe 2′,7′-dichlorodihydrofluorescein diacetate (DCFH-DA; Solarbio, Beijing, China) as described previously (32). Briefly, three independent cultures of *V. parahaemolyticus* YDE17 were grown to an OD_600_ of approximately 0.5. Cells were collected by centrifugation, washed twice with sterilized seawater, and resuspended to an OD_600_ of 0.5. Each suspension was divided into two equal aliquots: one was treated with 5 mM L-Cys for 30 min, and the other served as an untreated control. Subsequently, 1 μL of 10 mM DCFH-DA was added to each of the six samples, followed by incubation at 37 °C in the dark for 15 min. After washing, fluorescence intensity was measured using a multifunctional microplate reader (Varioskan Lux; Thermo Fisher Scientific, Shanghai, China) with excitation at 488 nm and emission at 525 nm. To confirm the role of ROS in L-Cys-induced cell death, bacterial cells were pretreated with 0.3% thiourea (a ROS scavenger) prior to L-Cys exposure, and fluorescence intensity was measured after 2 h of incubation.

### Bacterial Viability Assay

Bacterial viability was assessed using the LIVE/DEAD Bacterial Viability Kit (Beyotime, Shanghai, China) according to the manufacturer’s instructions. Briefly, *V. parahaemolyticus* YDE17 was grown to an OD_600_ of 0.5, washed twice with sterilized seawater, and resuspended to the same OD_600_. For treatment, bacterial suspensions were exposed to L-Cys at concentrations of 0, 1.25, 2.5, 5, 10, or 20 mM, with three replicates per concentration, and incubated at 28 °C for 30 min. To evaluate the role of ROS in cell death, 0.3% thiourea was added to parallel samples in combination with L-Cys. After staining with PI, fluorescence intensity was measured using a Varioskan Lux multimode microplate reader (Thermo Fisher Scientific, USA) at an excitation wavelength of 525 nm and an emission wavelength of 617 nm.

A standard curve was made to quantify cell mortality. Exponentially grown *V. parahaemolyticus* YDE17 was divided into two groups: live cells (resuspended in sterilized seawater) and dead cells (fixed with pure ethanol at room temperature for 1 h). Both groups were washed and adjusted to an OD_600_ of 0.5. Serial mortality standards of 100%, 80%, 60%, 40%, 20%, 10%, and 0% were prepared by mixing appropriate volumes of live and dead cell suspensions, e.g., 100% standard: 1 mL dead cell suspension; 80% standard: 0.8 mL dead suspension plus 0.2 mL live suspension, etc. Each standard was prepared in triplicate and measured under the same fluorescence conditions. The resulting standard curve was used to calculate mortality rates in the treatment groups.

### Measurement of Antioxidant Enzyme Activity and Glutathione disulfide (GSSG) Concentration

*V. parahaemolyticus* YDE17 was cultured with or without L-Cys, with three independent biological replicates per condition. Cells were harvested from each replicate, washed three times with sterilized seawater, and processed for enzyme activity assays. The activities of glutathione reductase (GR), catalase (CAT), and superoxide dismutase (SOD) were determined using commercial kits. GR and CAT kits were from Solarbio (Beijing, China); SOD kit was from Njjcbio (Nanjing, China). GR activity was defined as the amount of enzyme oxidizing 1 μmol NADPH per minute per 10⁴ cells at 37 °C and pH 8.0. CAT activity was expressed as the amount of enzyme degrading 1 μmol H_2_O_2_ per minute per 10⁵ cells. One unit of SOD activity (U) was defined as the amount of enzyme causing 50% inhibition in the specified assay system.

Intracellular oxidized glutathione (GSSG) content was measured using a GSH/GSSG Assay Kit (Beyotime, Shanghai, China) following the manufacturer’s protocol. Briefly, standard curves were generated using GSSG at different incubation time points. An optimal incubation time was selected based on these curves. Samples were then prepared accordingly, and the absorbance at 412 nm was measured. GSSG concentrations were calculated based on the standard curve.

### Effect of L-Cys on the Growth of Different *Vibrio* spp. Strains

To assess the ubiquity of L-Cys inhibition on *Vibrio* species, six strains, i.e., *V. alginolyticus* H1, *V. splendidus* SSD10, *V. anguillarum* SSD9, *V. mediterranei* RPV2, *V. harveyi* W18, and *V. crassostreae* S2, were cultured in the presence of L-Cys. The compound was added to M9 minimal medium at concentrations of 0, 1.25, 2.5, 5, 10, 20, 30, 40, and 50 mM, respectively. Each concentration was tested in triplicate using 15 mL glass flasks, yielding a total of 27 experimental samples per strain.

Additionally, M9 medium supplemented with 40 mM L-Cys was used to evaluate whether *Vibrio* spp. could utilize L-Cys as the sole carbon source. All cultures were incubated at 28 °C with continuous shaking for 12 h, after which the optical density at OD_600_ was measured.

### Statistical Analysis

Data are presented as mean ± SD. Statistical analysis was performed using GraphPad Prism software (GraphPad Software), with group comparisons conducted by one-way analysis of variance (ANOVA) and independent samples *t*-test. Statistical significance was defined as \**P* < 0.05, \*\**P* < 0.01; the labels “a” and “b” also denote *P* < 0.05 and were used to distinguish significance levels between different comparison groups. The raw sequencing data have been deposited in the NCBI SRA database under accession number PRJNA1394195.

## Results

### The effects of amino acids on the growth of *V. parahaemolyticus* YDE17

When individual amino acids were supplied as the sole carbon source on solid M9 agar plates, *V. parahaemolyticus* YDE17 exhibited distinct growth phenotypes, with noticeable differences in colony size and number across the 20 amino acids tested, indicating their varying efficiency as carbon substrates (**Supplementary Fig. 1**). In liquid medium with L-Ile or L-Cys as the sole carbon source, OD_600_ readings remained negligible after 48 h of culture, suggesting that these two amino acids are likely not used as carbon sources by *V. parahaemolyticus* YDE17 (**Fig. 1A**). Similarly, on solid medium with L-Cys as the sole carbon source, no visible colonies formed (**Fig. 1B**). All other amino acids supported growth to some extent; however, compared to colonies grown on nutrient-rich 2216E plates, colony numbers were reduced for every single amino acid tested. For example, when L-Ile and L-Met were individually provided as the sole carbon source, colony counts decreased by 79% and 57%, respectively (**Fig. 1B**). Together, these results indicate that among the 20 amino acids examined, L-Cys, unlike the others, cannot, or is only minimally able to, serve as a sole carbon source to support the growth of *V. parahaemolyticus* YDE17.

**Fig. 1.**
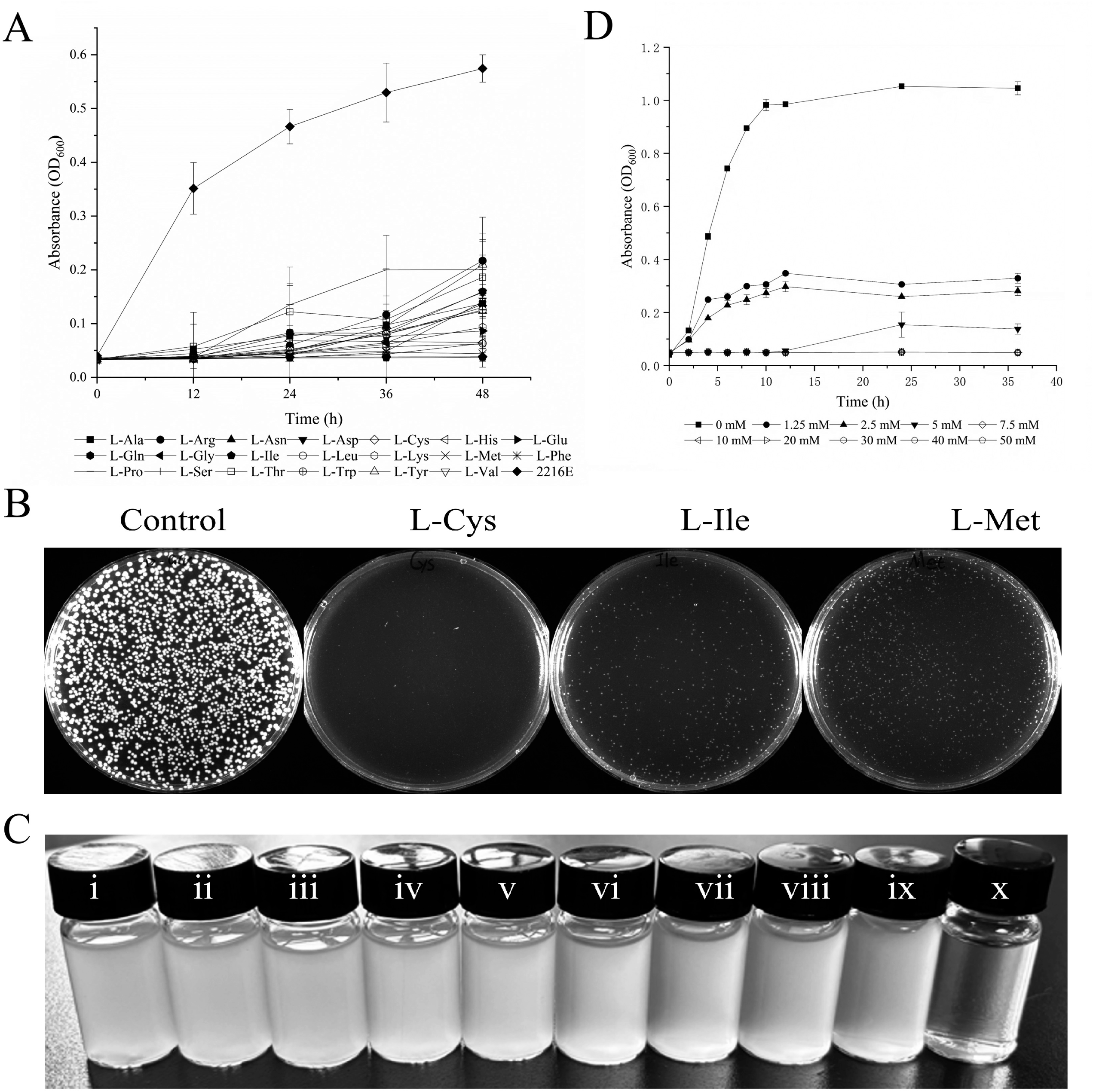
The growth of *V. parahaemolyticus* YDE17 in the presence of L-Cys under various conditions. (A) Growth of *V. parahaemolyticus* YDE17 in M9 medium supplemented with a single amino acid as the sole carbon source. Data are presented as mean ± SD. (B) Colony formation of *V. parahaemolyticus* YDE17 on M9 agar plates containing D-glucose (control), L-Cys, L-Ile, or L-Met as the sole carbon source. (C) Growth of *V. parahaemolyticus* YDE17 in M9 minimal medium supplemented with increasing concentrations of L-Cys. Symbols ⅰ–ⅸ correspond to L-Cys concentrations of 0, 1.25, 2.5, 5, 10, 20, 30, 40, and 50 mM, respectively; symbol x denotes M9 medium with 20 mM L-Cys as the sole carbon source, which did not support bacterial growth. (D) Growth of *V. parahaemolyticus* YDE17 in 2216E medium supplemented with different concentrations of L-Cys. Experiments were performed with three independent replicates for each concentration. Data are shown as mean ± SD.

### L-Cys inhibited the growth of *V. parahaemolyticus* YDE17

When varying concentrations of L-Cys were introduced into M9 minimal medium, the OD_600_ of *V. parahaemolyticus* YDE17 declined progressively with increasing L-Cys levels. No growth was observed when 40 mM L-Cys was supplied as the sole carbon source (**Fig. 1C**). These results suggested that L-Cys exerted a dose-dependent inhibitory effect on bacterial growth under these conditions. Although the inhibitory effect still present, L-Cys did not completely suppress growth even at 50 mM in M9 medium (**Fig. 1C**). In contrast, when cultured in 2216E medium supplemented with 0-50 mM L-Cys, a dose-dependent inhibition was evident within the range of 0-5 mM, leading to a gradual decrease in OD_600_ over 20 h. At concentrations above 7.5 mM, growth of *V. parahaemolyticus* YDE17 was completely inhibited (**Fig. 1D**).

### L-Cys Damaged Cellular Structure and Membrane Integrity

Scanning electron microscopy (SEM) revealed structural alterations in *V. parahaemolyticus* YDE17 cells following L-Cys treatment. Untreated cells exhibited typical short-rod morphology, abundant filamentous extracellular secretions, and intact cell surfaces. In contrast, L-Cys-treated cells displayed a pronounced reduction in filamentous secretions, accompanied by the appearance of numerous irregularly shaped and unevenly distributed pores in the cell wall (**Fig. 2A**). To further assess cellular damage, nucleic acid content in the culture supernatant was measured with increasing L-Cys concentrations. Higher L-Cys concentrations resulted in greater nucleic acid release into the supernatant (**Fig. 2B**), indicating L-Cys-induced disruption of intracellular integrity and leakage of cytoplasmic contents. Notably, significant membrane damage was observed within 15 min after exposure, but prolonging treatment time did not intensify this effect. Bacterial viability under L-Cys exposure was evaluated using fluorescence staining. In the untreated control, DMAO-stained live cells substantially outnumbered PI-stained dead cells. However, treatment with 5 mM L-Cys markedly altered this ratio, with a significant increase in PI-positive dead cells (**Fig. 2C**). These results demonstrate that 10 mM L-Cys directly induced cell death in *V. parahaemolyticus* YDE17.

**Fig. 2.**
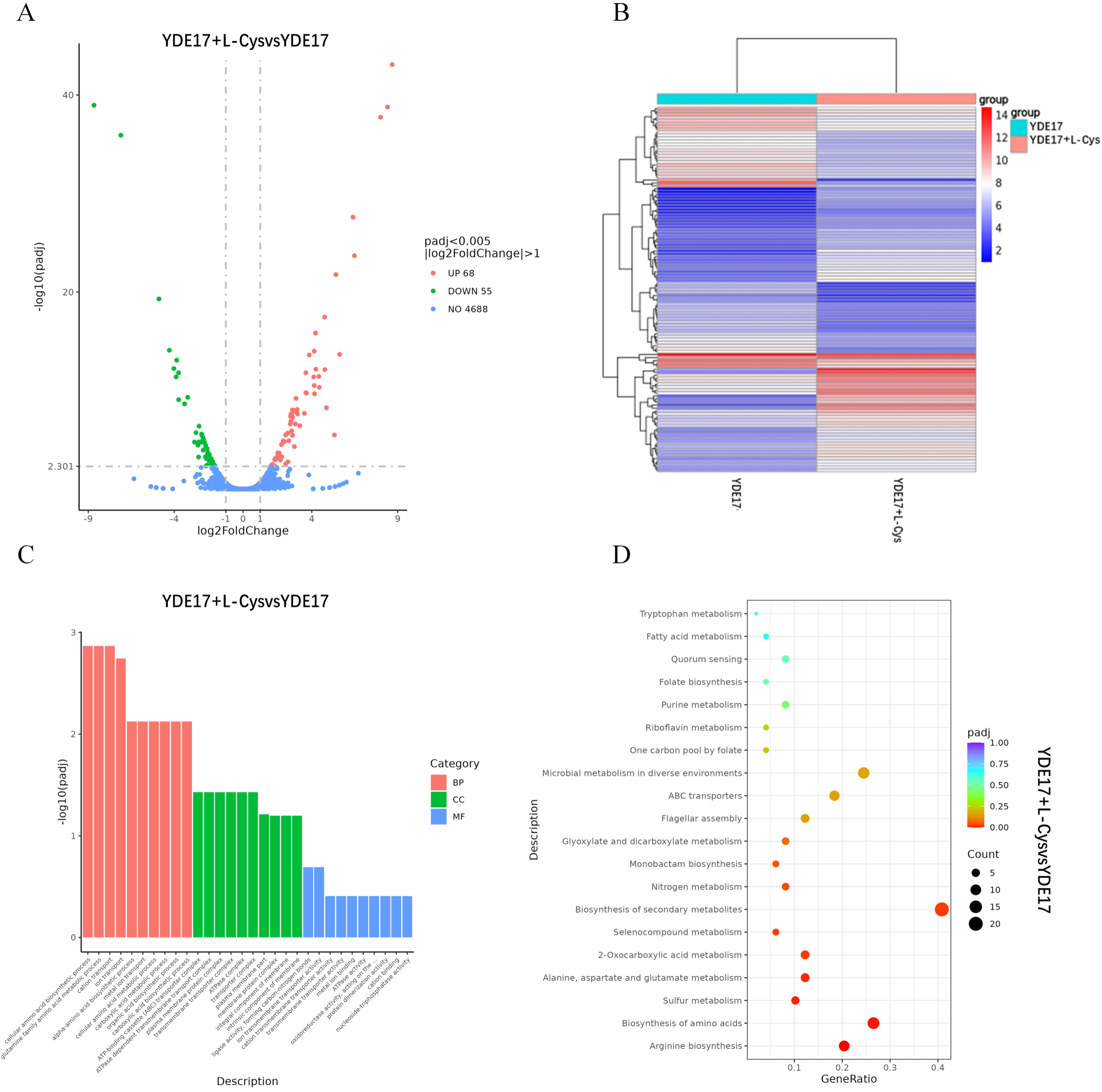
Phenotypic alterations in *V. parahaemolyticus* YDE17 cells following L-Cys exposure. (A) Scanning electron microscopy (SEM) images of individual *V. parahaemolyticus* YDE17 cells, untreated or treated with L-Cys. Arrows Ⅰ–Ⅵ highlight representative regions displaying compromised cell wall integrity. (B) Dose-and time-dependent extracellular nucleic acid release from *V. parahaemolyticus* YDE17 induced by L-Cys. Data are presented as mean ± SD. At each time point, the group without L-Cys was used as the control. Statistical significance was assessed using t-tests, with at least three biological replicates per group (\**P* < 0.05, \*\**P* < 0.01). (C) Confocal laser scanning microscopy images of untreated cells and cells treated with 10 mM L-Cys. Green fluorescence (DMAO staining) indicates cells with intact membranes, whereas red fluorescence (PI staining) labels cells with damaged membranes. Scale bar = 10 μm.

Taken together, these findings indicated that L-Cys rapidly caused severe damage to the cell membrane of *V. parahaemolyticus* YDE17, ultimately leading to bacterial death.

### L-Cys Enhances Bactericidal Effects by Increasing ROS

Transcriptomic analysis identified 123 DEGs between the YDE17 control and YDE17 + L-Cys treatment groups, comprising 68 upregulated and 55 downregulated genes (**Fig. 3A**). Expression levels of these DEGs were significantly altered in L-Cys-treated cells relative to the untreated control (**Fig. 3B**). Upregulated DEGs were primarily enriched in the following Gene Ontology (GO) categories: Biological processes (BP): localization (GO:0051179) and amino acid metabolic processes (GO:0008652); Cellular components (CC): membrane components (GO:0044425), ABC transporter complexes (GO:0043190), and various organelles (GO:0043226); Molecular functions (MF): enzyme activity (GO:0016787), ion binding (GO:0043168), and transporter activity (GO:0022857) (**Fig. 3C; Supplementary Fig. 2A; Supplementary Table 1**). Downregulated DEGs were mainly associated with: BP: localization (GO:0051234), ion transport (GO:0006812), and oxidation–reduction processes (GO:0055114); CC: organelles (GO:0043228), cell projections (GO:0042995), and bacterial-type flagella (GO:0009288); MF: enzyme activity (GO:0016491), ion binding (GO:0043167), and transporter activity (GO:0015075) (**Fig. 3C; Supplementary Fig. 2B; Supplementary Table 1**).

**Fig. 3.**
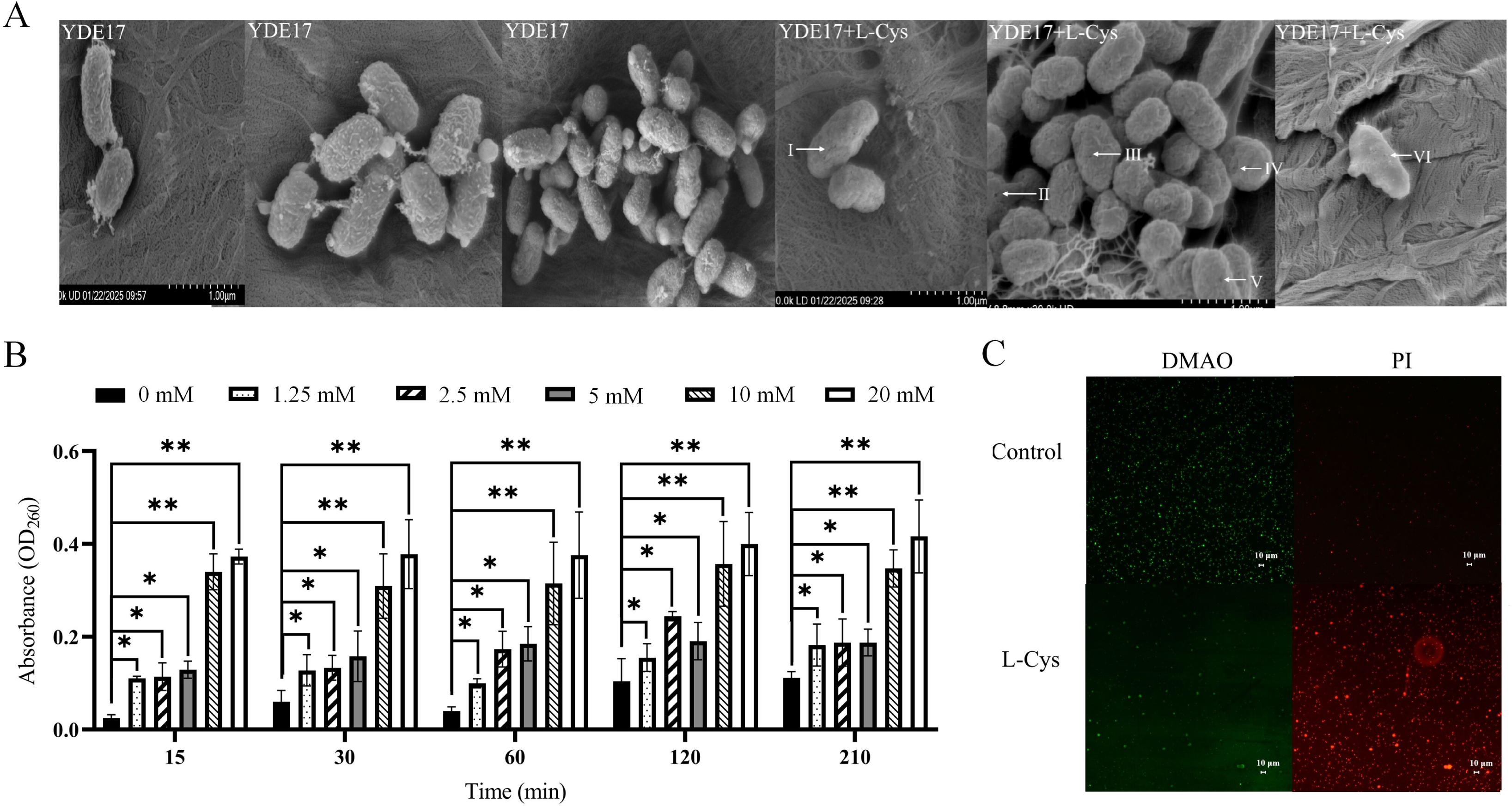
Transcriptomic analysis of *V. parahaemolyticus* YDE17 with and without L-Cys treatment. (A) Volcano plot of DEGs between YDE17 and YDE17 + L-Cys. The x-axis shows log_2_(fold change), and the y-axis represents −log₁₀(adjusted *P*-value). Red and green dots denote significantly upregulated and downregulated genes, respectively. (B) Hierarchical clustering of DEGs. Each row corresponds to an individual gene. Transcriptomic profiles between the two groups were visualized using MatLab. (C) GO enrichment analysis. The x-axis lists GO terms; the y-axis shows −log₁₀(adjusted *P*-value) to indicate enrichment significance. Bars represent significantly enriched terms in biological process (BP), cellular component (CC), and molecular function (MF) categories. (D) KEGG pathway enrichment analysis. The x-axis represents the ratio of DEG number to total gene number in a pathway; the y-axis lists pathway names. Point color and size correspond to the adjusted P-value and the percentage of DEGs in the pathway, respectively.

The term “Localization” (GO:0051179) emerged as the most enriched category, comprising 21 genes (42.9% of DEGs) and exhibiting bidirectional regulation. Upregulated genes were primarily associated with substrate transport and membrane-localization systems, whereas cation efflux pumps and ABC transporter subunits were downregulated, suggesting a broad activation of membrane transport functions. “Cellular amino acid biosynthetic process” (GO:0008652) showed the highest statistical significance (*P* = 1.77 × 10⁻⁵), with all seven related genes being upregulated, indicating its role as a central node in the amino acid metabolic network that is directly linked to the arginine biosynthesis pathway in KEGG. Seven DEGs were enriched in “oxidation–reduction process” (GO:0055114), among which *VP_RS14070*, *fadE*, *VP_RS14130*, and *VP_RS13375* were significantly downregulated (*P* < 0.05). Of particular note, *VP_RS14070*, which encodes superoxide dismutase, was markedly downregulated. Given its involvement in oxidation–reduction processes, metal ion binding, and antioxidant activity, we subsequently measured the intracellular ROS level in the following experiments.

KEGG pathway analysis revealed that DEGs were enriched in several metabolic and transport categories, including biosynthesis of secondary metabolites, amino acid biosynthesis, arginine biosynthesis, microbial metabolism in diverse environments, and ABC transporters (**Fig. 3D; Supplementary Figs. 2C, D; Supplementary Table 1**). Among the upregulated pathways, arginine biosynthesis (vpa00220), riboflavin metabolism (vpa00740), and fatty acid biosynthesis (vpa00061) were notably enriched. Arginine biosynthesis emerged as the most statistically significant term (*P* = 5.30 × 10⁻¹¹) and was also the most highly enriched in both KEGG and GO analyses. This pathway contained ten genes, *argC*, *argB*, *argD*, *argF*, *argG*, *argH*, *carA*, *carB*, *astA*, and *astB*, all of which were upregulated. In contrast, pathways such as one-carbon pool by folate (vpa00670), sulfur metabolism (vpa00920), and flagellar assembly (vpa02040) were downregulated. Sulfur metabolism showed the strongest downregulation, with all five related genes, *cysI*, *cysJ*, *cysG*, *sir*, and *phsA*, being suppressed.

These results indicate that L-Cys substantially alters the physiological state of *V. parahaemolyticus* by modulating multiple key metabolic and biosynthetic pathways.

### L-Cys Elevated ROS through Hydrogen Sulfide (H_2_S) Production

Production of H_2_S by *V. parahaemolyticus* YDE17 in the presence of exogenous L-Cys was assessed both qualitatively and quantitatively. Qualitatively, a distinct rotten-egg odor was noted during bacterial culture, and blackening of lead acetate test strips confirmed H_2_S release. The darkness of the test strips increased progressively as L-Cys concentration rose from 0 to 2.5 mM. At L-Cys concentrations above 2.5 mM, the strips turned completely black, indicating that at lower concentrations, H_2_S levels depended on the supplied amount of L-Cys (**Fig. 4A**). Quantitative measurements showed that after 4 h of incubation, H_2_S levels reached 60 ± 8 ppm with 1.25 mM L-Cys, 126.67 ± 20.13 ppm with 2.5 mM L-Cys, and 252.33 ± 12.58 ppm with 5 mM L-Cys (**Fig. 4B**).

**Fig. 4.**
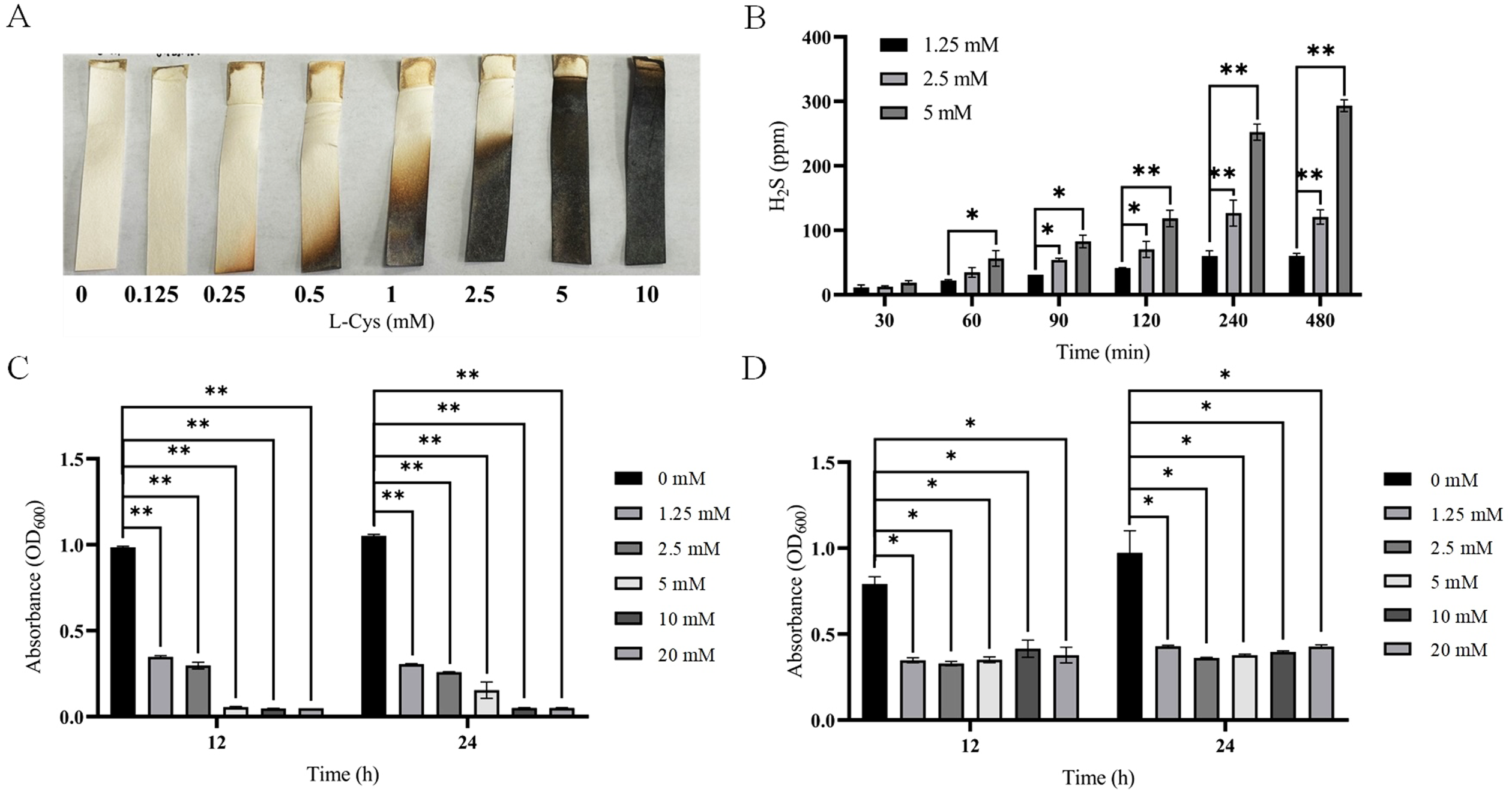
H_2_S production by *V. parahaemolyticus* YDE17 in the presence of L-Cys contributes to growth inhibition. (A) Qualitative detection of H_2_S using lead acetate test paper. Darkening of the test paper with increasing L-Cys concentration indicates elevated H_2_S generation. (B) Quantitative measurement of H_2_S levels. (C) Growth of *V. parahaemolyticus* YDE17 in 2216E medium supplemented with L-Cys, as measured by OD_600_. (D) OD_600_ of *V. parahaemolyticus* YDE17 cultured in 2216E medium with NaHS as an exogenous H_2_S donor. All data are presented as mean ± SD. For each assay, medium without L-Cys served as the control. Statistical significance was evaluated by t-test, with at least three biological replicates per group (\**P* < 0.05, \*\**P* < 0.01).

Increasing L-Cys concentrations resulted in elevated H_2_S accumulation in a dose- and time-dependent manner; however, concentrations above 10 mM completely suppressed bacterial growth (**Fig. 4C**). To clarify whether L-Cys acts primarily through H_2_S, NaHS was chosen as an exogenous H_2_S donor. The addition of NaHS also inhibited the growth of *V. parahaemolyticus* YDE17 in a dose-dependent fashion, although its inhibitory effect was notably weaker than that of L-Cys at equivalent concentrations. Below 5 mM, both compounds showed similar inhibitory activity, with OD_600_ gradually declining as concentration increased. Above 5 mM, however, L-Cys completely suppressed growth, while the inhibitory effect of NaHS plateaued and failed to achieve complete growth arrest even at higher concentrations (**Fig. 4C, D**).

Based on transcriptomic findings, we evaluated antioxidant-related enzyme activities in *V. parahaemolyticus* YDE17 following L-Cys treatment by measuring key antioxidant enzymes (SOD, CAT, and GR) and GSSG content. After exposure to 5 mM L-Cys, SOD activity decreased by 82.71% (**Fig. 5A**), while GR activity was significantly reduced by 16.51% (**Fig. 5B**). CAT activity remained unchanged (**Fig. 5C**). Concurrently, GSSG levels increased from 1.561 μM to 2.898 μM, representing an 85.6% rise (**Fig. 5D**). These results indicate that L-Cys disrupts cellular redox homeostasis by impairing major antioxidant defenses and disturbing glutathione system regulation.

**Fig. 5.**
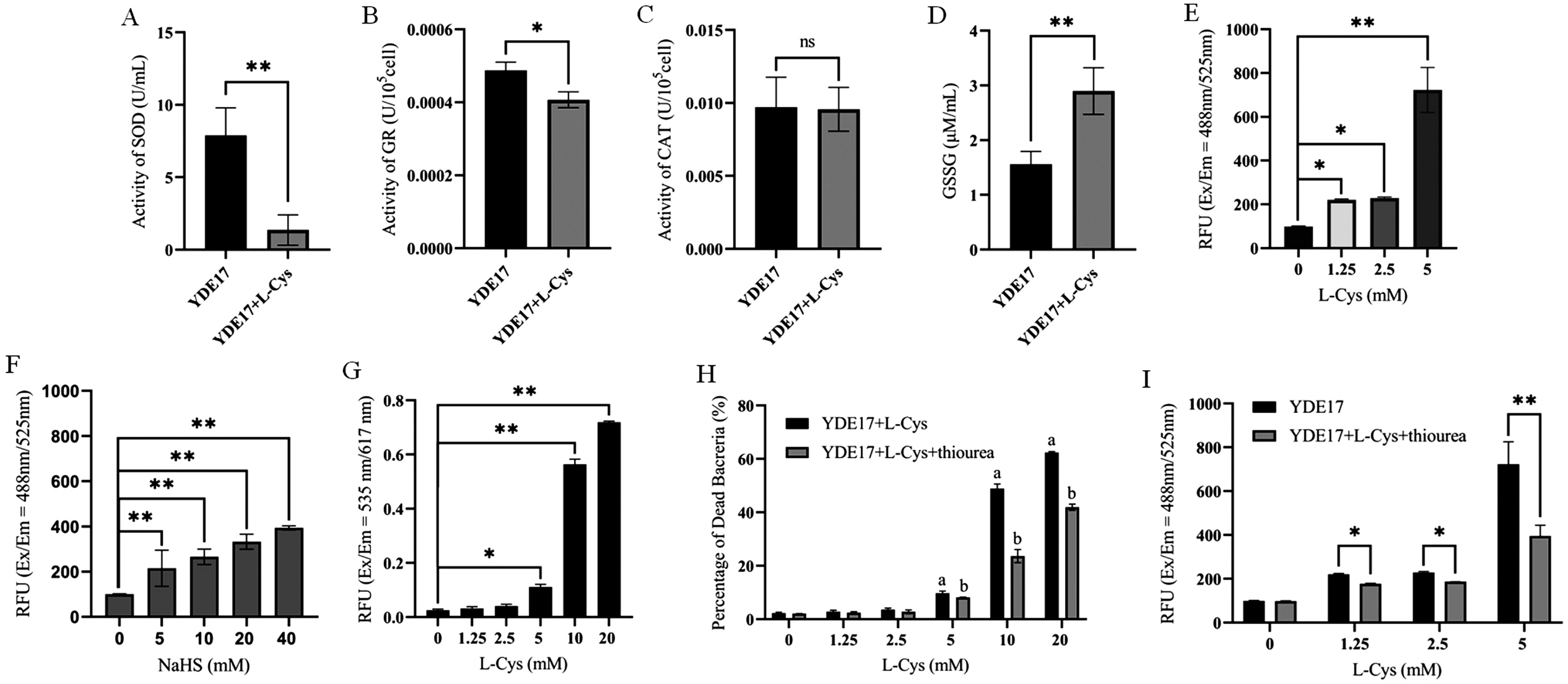
L-Cys treatment reduced antioxidant enzyme activity while inducing oxidative stress and membrane damage in *V. parahaemolyticus* YDE17. (A) SOD activity in *V. parahaemolyticus* YDE17 with or without L-Cys. (B) GR activity with or without L-Cys. (C) CAT activity with or without L-Cys. (D) Intracellular GSSG levels with or without L-Cys. (E) Effect of L-Cys on intracellular ROS levels. (F) Effect of H_2_S on intracellular ROS levels. (G) L-Cys-induced membrane permeability assessed by PI staining. (H) Mortality rate of *V. parahaemolyticus* YDE17 treated with different concentrations of L-Cys, and the intervening effect of thiourea. Black bars represent mortality in the “YDE17 + L-Cys” group; gray bars represent mortality in the “YDE17 + L-Cys + thiourea (0.3%)” group. Letters a and b denote significant inter-group differences (*P* < 0.05): a indicates comparison against the 0 mM L-Cys control; b indicates comparison between the thiourea-supplemented group and its corresponding L-Cys-only group at the same concentration. (I) Effect of the ROS scavenger thiourea on ROS production. ROS levels under L-Cys, NaHS, and thiourea treatments were measured using a multi-mode microplate reader (Ex/Em = 488 nm/525 nm). Membrane permeability changes were assessed using the same reader (Ex/Em = 535 nm/617 nm). All values are expressed as mean ± SD from three independent experiments. The group without L-Cys served as the control in all assays, and significance was determined by *t*-test (\**P* < 0.05, \*\**P* < 0.01).

Based on the above results, the intracellular ROS levels in the presence of L-Cys was measured. As shown in **Fig. 5E**, ROS fluorescence intensity increased from 98.8 RFU in untreated controls to 220.2 ± 4.10 RFU in the presence of 1.25 mM L-Cys, 228.6 ± 5.52 RFU in the presence of 2.5 mM L-Cys, and 722.6 ± 102.89 RFU in the presence of 5 mM L-Cys, demonstrating that L-Cys significantly induced ROS production and acts as a potent pro-oxidant. To dissect the contributions of L-Cys and its metabolite H_2_S, parallel experiments with NaHS an H_2_S donor were performed. ROS fluorescence intensity in the presence of 10 mM NaHS reached only 265.8 ± 33.7 RFU, which was markedly lower than that 722.6 ± 102.89 RFU observed with an equivalent concentration of L-Cys (**Fig. 5F**). This suggests that beyond the H_2_S pathway, L-Cys likely elevated ROS levels through additional mechanisms.

Further membrane integrity assessment via PI staining showed that L-Cys concentrations above 5 mM significantly increased membrane permeability in *V. parahaemolyticus* YDE17 (**Fig. 5G**). Based on a bacterial viability standard curve (**Supplementary Fig. 3B**), mortality rates reached approximately 9.68 ± 0.83%, 48.91 ± 1.64%, and 62.36 ± 0.40% in groups treated with 5 mM, 10 mM, and 20 mM L-Cys, respectively (**Fig. 5H**). These results indicated that L-Cys-induced ROS accumulation led to structural damage of the bacterial membrane, ultimately causing irreversible cell death.

To determine whether ROS directly contributed to bacterial death, the ROS scavenger thiourea, at a concentration of 0.3 was added to the culture. As shown in **Fig. 5I**, thiourea significantly attenuated the ROS levels induced by L-Cys. Correspondingly, PI staining revealed that thiourea also markedly improved bacterial survival under L-Cys treatment (**Fig. 5H**). Together, these data confirmed that the ROS inhibitor thiourea reduced both intracellular ROS and L-Cys-induced mortality in *V. parahaemolyticus* YDE17, supporting the conclusion that excessive ROS accumulation directly mediated bacterial cell death.

### L-Cys Inhibits the Swimming Motility of *V. parahaemolyticus* YDE17

We further investigated whether L-Cys influences virulence-associated traits in *V. parahaemolyticus* YDE17 beyond its growth inhibitory effect. In soft-agar swimming assays, untreated bacteria formed an approximately spherical migration zone with a diameter of 2.07 ± 0.27 cm, with cell density gradually decreasing from the center outward. In contrast, in the presence of 5 mM L-Cys, migration was largely confined near the inoculation site, resulting in a significantly smaller migration zone (1.32 ± 0.08 cm; **Fig. 6A**), indicating that L-Cys inhibits swimming motility. For swarming motility, the control group formed a migration zone of approximately 0.7 ± 0.01 cm in diameter, while the L-Cys-treated group showed a slightly larger zone (0.9 ± 0.03 cm). The difference was not statistically significant, suggesting that L-Cys does not substantially affect swarming motility (**Fig. 6B**).

**Fig. 6.**
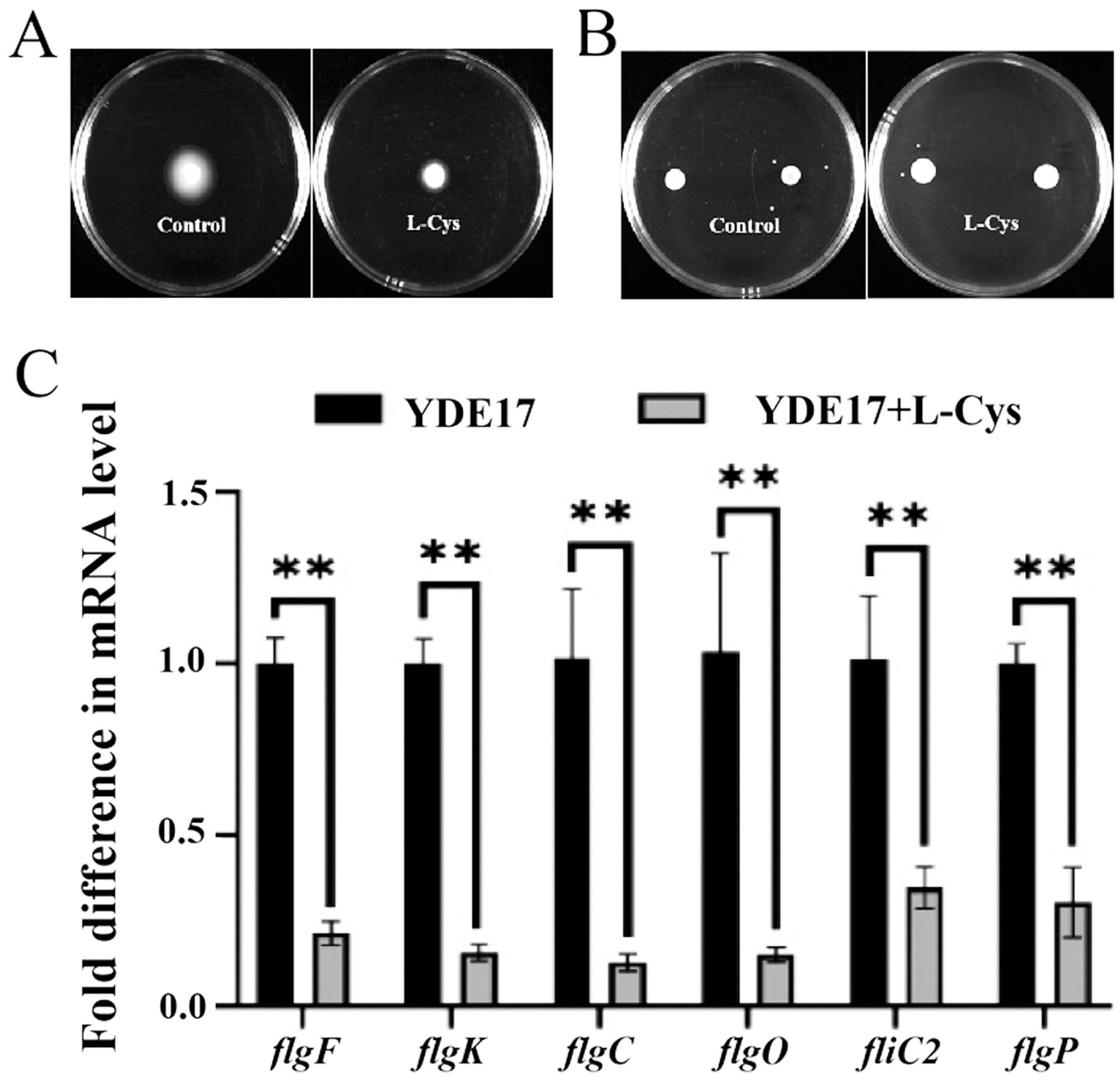
Motility alterations in *V. parahaemolyticus* YDE17 induced by exogenous L-Cys. (A) Swimming assay in 2216E soft agar. Control group: 0 mM L-Cys; treatment group: 10 mM L-Cys. Plates were inoculated with 2 µL of bacterial suspension and incubated statically at 28 °C for 12 h before imaging. (B) Swarming assay in 2216E semi-solid agar. Control: 0 mM L-Cys; treatment: 10 mM L-Cys. Plates were inoculated with 5 µL of bacterial suspension and incubated statically at 28 °C for 12 h before imaging. (C) qRT-PCR analysis of flagellar assembly related genes identified via KEGG pathway mapping. Gene expression levels in L-Cys-treated cells (YDE17 + L-Cys) were compared to the untreated control (YDE17). Data represent three independent experiments; significance was assessed by t-test (\*\**P* < 0.01).

These phenotypic observations aligned with transcriptomic data, which showed that genes involved in flagellar assembly, including *flgO*, *flgP*, *flgK*, *flgF*, *fliC*, and *fliC2*, were downregulated after 5 mM L-Cys treatment. This downregulation was further confirmed by real-time RT-PCR (**Fig. 6C**). The reduced expression of these flagellar-related genes likely impairs the function of key flagellar structures, such as the H ring, rod, L/P ring, hook-filament junction, and filament, thereby diminishing bacterial motility.

### L-Cys Exhibited a Universal Inhibitory Effect on *Vibrio* Growth

We further investigated whether L-Cys exhibits broad-spectrum inhibitory activity against diverse marine *Vibrio* species. Six *Vibrio* strains, *V. alginolyticus* H1, *V. splendidus* SSD10, *V. anguillarum* SSD9, *V. mediterranei* RPV2, *V. harveyi* W18, and *V. crassostreae* S2, were tested for growth in the presence of L-Cys at concentrations ranging from 1.25 to 50 mM. All six strains showed inhibited growth under these conditions. Notably, *V. anguillarum* SSD9 was completely inhibited by L-Cys, whereas *V. alginolyticus* H1, *V. splendidus* SSD10, *V. mediterranei* RPV2, *V. harveyi* W18, and *V. crassostreae* S2 displayed a dose-dependent inhibition pattern similar to that observed in *V. parahaemolyticus* YDE17, with progressively stronger suppression as L-Cys concentration increased (**Fig. S3C**).

## Discussion

In this study, we demonstrated for the first time that *V. parahaemolyticus* is highly sensitive to L-Cys and unable to use it as the sole carbon source. The addition of L-Cys to the culture medium significantly inhibited the growth of *V. parahaemolyticus* YDE17 in a dose-dependent manner. This growth suppression was directly linked to the disruption of cell membrane integrity, consistent with the previously reported antibacterial activity of L-Cys against *E. coli* (24, 34), *S. aureus* (24), and *Monilinia fructicola* (25). Notably, L-Cys exhibits a dual, concentration-dependent effect in *E. coli*: at concentrations lower than 1 mM, it serves as a sulfur source and reducing agent that alleviates oxidative stress and promotes growth (35, 36); whereas at concentrations higher than 5 mM, it inhibits growth through multiple mechanisms, including metabolic interference (37). However, such a dual effect was not observed in *V. parahaemolyticus* YDE17. This difference may be due to the relatively high L-Cys concentrations used in our study, or to intrinsic strain-specific traits and medium composition. We observed that L-Cys concentrations above 7.5 mM completely inhibited the growth of *V. parahaemolyticus* YDE17 in 2216E medium, whereas even 100 mM L-Cys did not achieve full suppression in M9 minimal medium. Together, these findings suggested that the impact of exogenous L-Cys on microorganisms is not only concentration- and species-dependent, but also strongly influenced by the nutritional composition of the medium. This phenomenon may be closely related to microbial metabolic networks, antioxidant capacity, and L-Cys transport efficiency (24), as has similarly been reported in fungal pathogens (25).

Previous studies indicate that L-Cys exerts antibacterial effects through multiple pathways, including ROS generation via auto-oxidation to induce oxidative damage (32), and H_2_S production that disrupts the electron transport chain (32, 38). In *E. coli*, however, existing work has often focused on one specific aspect such as H_2_S-mediated oxidative damage (32), synergistic bactericidal effects with hydrogen peroxide (39), or metabolic interference-driven growth inhibition (40). These studies remain largely confined to membrane level effects, without further exploration of oxidative stress or metabolic responses. But, this study systematically integrated a synergistic antibacterial mechanism that encompassed “metabolic network interference H_2_S-regulated oxidative stress ROS-mediated oxidative damage,” offering a more comprehensive perspective for understanding the antibacterial activity of L-Cys.

Transcriptomic analysis further elucidated the molecular mechanisms underlying L-Cys-induced disruption of redox homeostasis in *V. parahaemolyticus* YDE17. DEGs were primarily enriched in processes including substance localization, amino acid metabolism, ion transport, and redox reactions. Among them, three key expression shifts synergistically drove the observed ROS burst: First, downregulation of the superoxide dismutase gene *VP_RS14070* directly compromised the bacterial ROS-scavenging capacity, consistent with previously reported decreases in SOD activity in *E. coli* (41). Second, suppression of redox-related processes (GO:0055114) and sulfur metabolism (vpa00920) jointly disrupted the antioxidant defense system, providing molecular support for ROS-mediated bacterial death (42). Third, upregulation of *ribA*, encoding a rate-limiting enzyme in riboflavin synthesis, likely led to accumulation of flavin mononucleotide (FMN) and flavin adenine dinucleotide (FAD), which can promote electron leakage from the respiratory chain and further stimulate ROS production (43). Together, these transcriptional changes induced an intracellular ROS burst, ultimately leading to oxidative damage and bacterial death.

Notably, the toxic mechanism of L-Cys extends beyond ROS-mediated oxidative damage. Transcriptomic analysis revealed a significant upregulation of key genes in the arginine biosynthesis pathway (*argB*, *argC*, *argD*, and *argH*), which not only promotes de novo arginine synthesis and expands the intracellular arginine pool (30), but also elevates pathway flux to supply critical substrates for downstream redox metabolism and electron transfer, thereby indirectly driving substantial electron flow (44). Concurrently, L-Cys broadly suppressed central metabolic and repair pathways in the bacteria. Downregulation of *FadE*, a key enzyme in fatty acid β-oxidation, disrupted lipid metabolism and energy production (45). Reduced expression of assimilatory sulfite reductase (VPA0803), glycine cleavage system T-protein (VPA0805), and ribonucleotide reductase (VPRS09190) impaired one-carbon metabolism, serine metabolism, and nucleotide synthesis. This extensive inhibition compromised the synthesis and repair of proteins, lipids, nucleotides, and DNA, thereby depriving cells of the capacity to counteract oxidative damage (46, 47). The combination of oxidative injury and metabolic-repair blockade likely explains why 5 mM L-Cys exhibits substantially stronger antibacterial activity than a single H_2_S donor such as NaHS.

Moreover, the potent inhibitory effect of L-Cys on *V. parahaemolyticus* YDE17 was likely closely related to its enzymatic conversion into high intracellular concentrations of H_2_S. H_2_S exhibits dual toxicity: direct toxicity by blocking the respiratory chain and inducing an energy crisis (38), and indirect toxicity via suppression of antioxidant enzymes and promotion of ROS accumulation, which together amplify oxidative damage (48). This is consistent with previous reports that elevated H_2_S can inhibit antioxidant enzyme activity (26), disrupt the electron transport chain (6), and induce ROS accumulation to suppress microbial growth (48). Notably, endogenous H_2_S produced from L-Cys metabolism showed stronger antibacterial activity than exogenous H_2_S donors such as NaHS. This difference may be attributed to spatiotemporal specificity: endogenous enzymatic generation enables targeted, high-concentration accumulation of H_2_S inside bacterial cells (49), whereas exogenous donors are subject to dilution and loss during diffusion, resulting in lower effective intracellular concentrations (26, 49). Such localized H_2_S production directly targets and impairs the respiratory chain and antioxidant system, thereby enhancing bacteriostatic efficacy.

Treatment with 5 mM L-Cys significantly compromised the antioxidant defense system of *V. parahaemolyticus* YDE17, as evidenced by marked reductions in SOD and GR activities. SOD eliminates superoxide anions, while GR sustains GSH regeneration; diminished activity of these enzymes led to GSH depletion, weakened ROS-scavenging capacity, and disruption of redox homeostasis. Consistent with this, we observed that L-Cys significantly promoted ROS accumulation, aligning with the established paradigm in which exogenous chemicals induce excessive ROS to mediate bacterial death (50). These findings are in agreement with earlier reports that L-Cys promotes ROS accumulation in *E. coli* B2 (40), and that H_2_S reduces SOD and GR activities in *E. coli*, leading to ROS imbalance and altered GSH levels that ultimately cause cellular damage or death (32). It is noteworthy that at the same concentration, L-Cys induced higher ROS levels than the H_2_S donor NaHS. This difference may be explained by two synergistic mechanisms: first, endogenous H_2_S generated from L-Cys creates an instantaneous high-concentration local microenvironment within *V. parahaemolyticus* YDE17, enhancing ROS induction through localized enrichment; second, L-Cys likely amplifies oxidative stress via additional metabolic pathways, such as downregulation of *fhuF*, a gene involved in iron-sulfur cluster homeostasis (33), which may disrupt cluster balance and exacerbate oxidative damage. This aligns with the classical concept that high intracellular cysteine can drive the Fenton reaction and promote oxidative DNA damage (51), further supporting the multi-mechanistic synergy underlying L-Cys-induced ROS accumulation.

The observed reduction in swimming motility in *V. parahaemolyticus* YDE17 supports the metabolic regulation of virulence, a phenomenon also noted in *Cryptococcus neoformans*, where overexpression of the RNA-binding protein Puf4 reprograms carbon metabolism and elevates ROS levels, significantly decreasing the synthesis of virulence factors such as capsule and melanin (52). Notably, we extended the investigation of L-Cys inhibition to multiple pathogenic *Vibrio* species. The supplementation of L-Cys suppressed the growth of all tested *Vibrio* strains, indicating that L-Cys may possess broad-spectrum inhibitory activity against pathogenic *Vibrios*. Collectively, the potent inhibitory efficacy and wide inhibitory spectrum of L-Cys provide a promising basis for development of strategies to control *V. parahaemolyticus* and related pathogens infections.

## Conclusion

This study identified L-Cys as a bacteriostatic agent and systematically elucidated its inhibition against *V. parahaemolyticus* YDE17 through a triple synergistic pathway. First, exogenously supplied L-Cys is metabolized to generate high concentrations of H_2_S within the bacterial cell. This endogenous accumulation directly overwhelms antioxidant defenses. Second, a burst of ROS is induced via multiple routes, producing strong oxidants that damage cellular integrity and DNA. Third, key metabolic pathways, including sulfur metabolism and nucleotide synthesis, are disrupted, impairing the capacity for self-repair of the cells. Together, these multi-targeted effects lead to an irreversible collapse of redox homeostasis, further compromise membrane structure and function, and ultimately cause cell death. The broad inhibitory activity of L-Cys was also confirmed across other *Vibrio* species, indicating its potential as a wide-spectrum bacteriostatic agent. These findings offer new perspectives for developing novel strategies to control *Vibrio* infections in aquaculture.

## Data Availability Statement

The data presented in this study are available from the corresponding author upon reasonable request.

## Author contributions

**Qiuyan Yang:** Conceptualization, methodology, software, formal analysis, validation, investigation, resources, writing-original draft, writing-review&editing.

**Huaipeng Fang:** methodology, investigation.

**Min Meng:** validation, investigation.

**Qingxi Han:** resources, writing-review and editing, funding acquisition.

**Jinlin Xu:** supervision, funding acquisition.

**Weiwei Zhang:** conceptualization, methodology, visualization, supervision, project administration, writing–review and editing, funding acquisition.

## Acknowledgment

This work was finally supported by the National Natural Science Foundation of China (42376103), the Major Project of Science, Technology and Innovation 2025 in Ningbo City (2021Z007), the earmarked fund for CARS-49, and was also partly sponsored by K.C. Wong Magna Fund in Ningbo University.

---

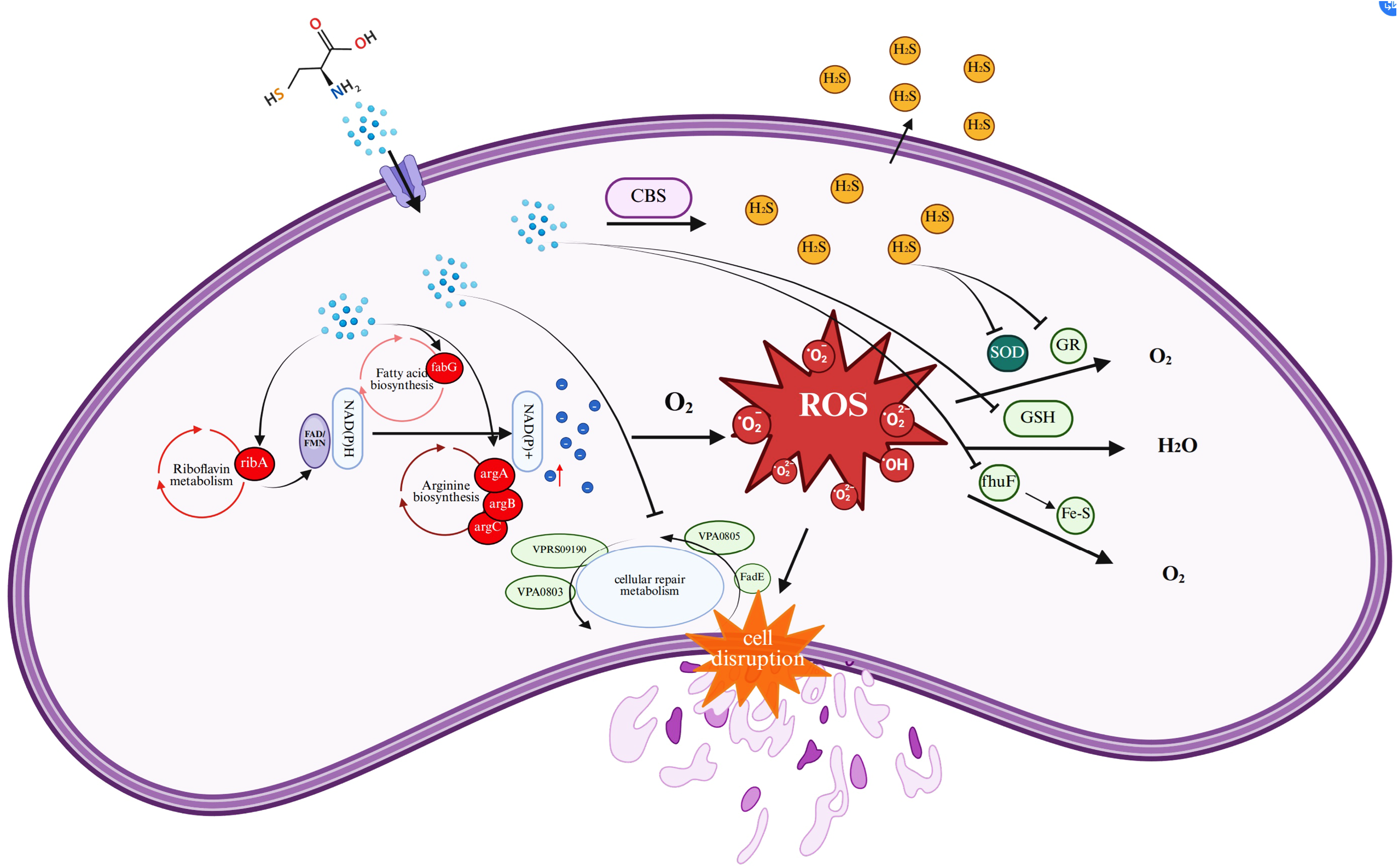

